# Distribution of haploid chromosomes into separate nuclei in two pathogenic fungi

**DOI:** 10.1101/2024.11.05.622059

**Authors:** Yan Xu, Lei Tian, Jinyi Tan, Josh Li, Nigel O’Neil, Martin Hirst, Phil Hieter, Yuelin Zhang, Xin Li

**Author notes:** These authors contributed equally to this work (Yan Xu, Lei Tian, Jinyi Tan).

## Abstract

The presence of nuclei defines eukaryotes, enabling compartmentalization of macromolecules and cellular regulation. Inside the nucleus, chromosome numbers vary greatly across organisms, both in terms of ploidy status and haploid chromosome number. For cells harboring multiple nuclei as in many fungal mycelia and animal muscle cells, each healthy nucleus is traditionally believed to carry at least one haploid set of chromosomes. Abnormal chromosome numbers in nuclei are often associated with aging, diseases such as cancer, developmental disorders or lethality. Here we report a surprising discovery that chromosomes in haploid cells of the fungal species *Sclerotinia sclerotiorum* and *Botrytis cinerea* are segregated into separate nuclei, with each nucleus containing only a fraction of the chromosomes. This is the first report of eukaryotic cells that partition chromosome subsets into distinct nuclei, bringing new questions and opening fresh avenues for chromosome biology.

---

*Sclerotinia sclerotiorum* (Lib.) de Bary (*S. sclerotiorum*) is one of the most notorious soilborne fungal pathogens. This ascomycete can colonize over 600 plant species (Liang and Rollins, 2018), causing severe yield and quality loss in many important crops including soybean and canola (Hegedus and Rimmer, 2005; Bolton, Thomma and Nelson, 2006). During its life cycle, *S. sclerotiorum* produces a melanized structure termed sclerotia, which can remain alive in soil for years (Adams and Ayers, 1979; Willetts and Wong, 1980). Depending on environmental conditions, sclerotia can germinate either as hyphae or release ascospores through a reproductive cup-shaped apothecia fruiting body to infect adjacent plants (Bolton, Thomma and Nelson, 2006). Another ascomycete *Botrytis cinerea* is the most damaging greenhouse pathogen. The massive amount of airborne haploid conidia enables its fast spread, while it can also form sclerotia to overwinter. Although both pathogens are economically important, the molecular mechanisms of their basic biology such as pathogenesis processes and sclerotia formation are poorly understood, partly due to the multinucleated nature of their mycelial cells hindering efficient genetic dissection (Ford *et al*., 1995).

To study the sclerotia formation process, we designed a forward genetic screen using ascospores of *S. sclerotiorum* (Xu *et al*., 2022). Pulsed-field gel electrophoresis based karyotyping and *de novo* genome assembly revealed that this ascomycete has 16 chromosomes (1N=16) (Fraissinet-Tachet, Reymond-Cotton and Fèvre, 1996; Amselem *et al*., 2011). As each ascospore has two nuclei, which are thought to be derived from mitotic division following meiosis (Ekins, 1999), we anticipated that each nucleus would carry the full 16 chromosomes. The binucleated ascospore should contain two identical haploid nuclei, resembling a diploid cell. Thus, after UV induced random mutagenesis morphological mutants should grow out as half sectors in the colony that originated from each ascospore. However, among over 80 nonsclerotial mutants isolated, we did not observe any sectoring patterns in their colonies (Xu *et al*., 2022 and unpublished data). In addition, if each nucleus within the binucleated ascospore carries a full set of chromosomes, a majority of UV-induced mutations should exist in a “pseudo-heterozygous” state with an approximate 1:1 ratio of mutation: wild-type in the resulting haploid mycelial colony. However, in all the mutants, including some non-morphological mutants we sequenced the mutations were homozygous with next-generation sequencing, most mutation reads did not show a “pseudo-heterozygous” state, with almost 100% mutation without the wild-type allele (Xu *et al*., 2022; Supplementary Fig. S1), arguing against the heterozygosity of the two nuclei. Our data rather suggest that each ascospore behaves genetically as a haploid cell. To us, the most straight forward explanation for such observation could be that each ascospore carries a full chromosome complement divided into the two nuclei instead of each nucleus containing all 16 chromosomes. Under such model, the ascospore is haploid with the full set of 16 chromosomes separated across two nuclei. As such hypothesis is against the established chromosome biology principle, we started examining the nuclei and chromosomes of *S. sclerotiorum*.

## Each ascospore of *S. sclerotiorum* contains two nuclei, likely with 8 chromosomes in each nucleus

To generate the ascospores, apothecia were induced from sclerotia of wild-type *S. sclerotiorum* strain 1980 (Fig. 1A). After staining with DAPI (4’,6-diamidino-2-phenylindole), which stains DNA, the ascospores were examined under a fluorescence microscope. As shown in Fig 1B, each ascospore indeed contains two DAPI-stained signals, likely representing two nuclei. To confirm the DAPI signals are truly from two nuclei, transmission electron microscopy (TEM) was carried out on ascospores. Indeed, two clear nuclei and nucleolus were observed with nucleolus present (Fig. 1C). As ascomycete nuclei are less than 5 micrometers in diameter, the DAPI-stained entities most likely representing chromosomes are only visible after the cell and nuclear envelope membranes are dissolved during fixation on glass slides. After scanning hundreds of broken ascospores, we were able to conclude that chromosome number in each ascospore is closer to 16 rather than 32 (a representative broken ascospore is shown in Fig. 1D), supporting our hypothesis that each ascospore, not each nucleus, contains a full haploid chromosome complement.

**Figure 1.**
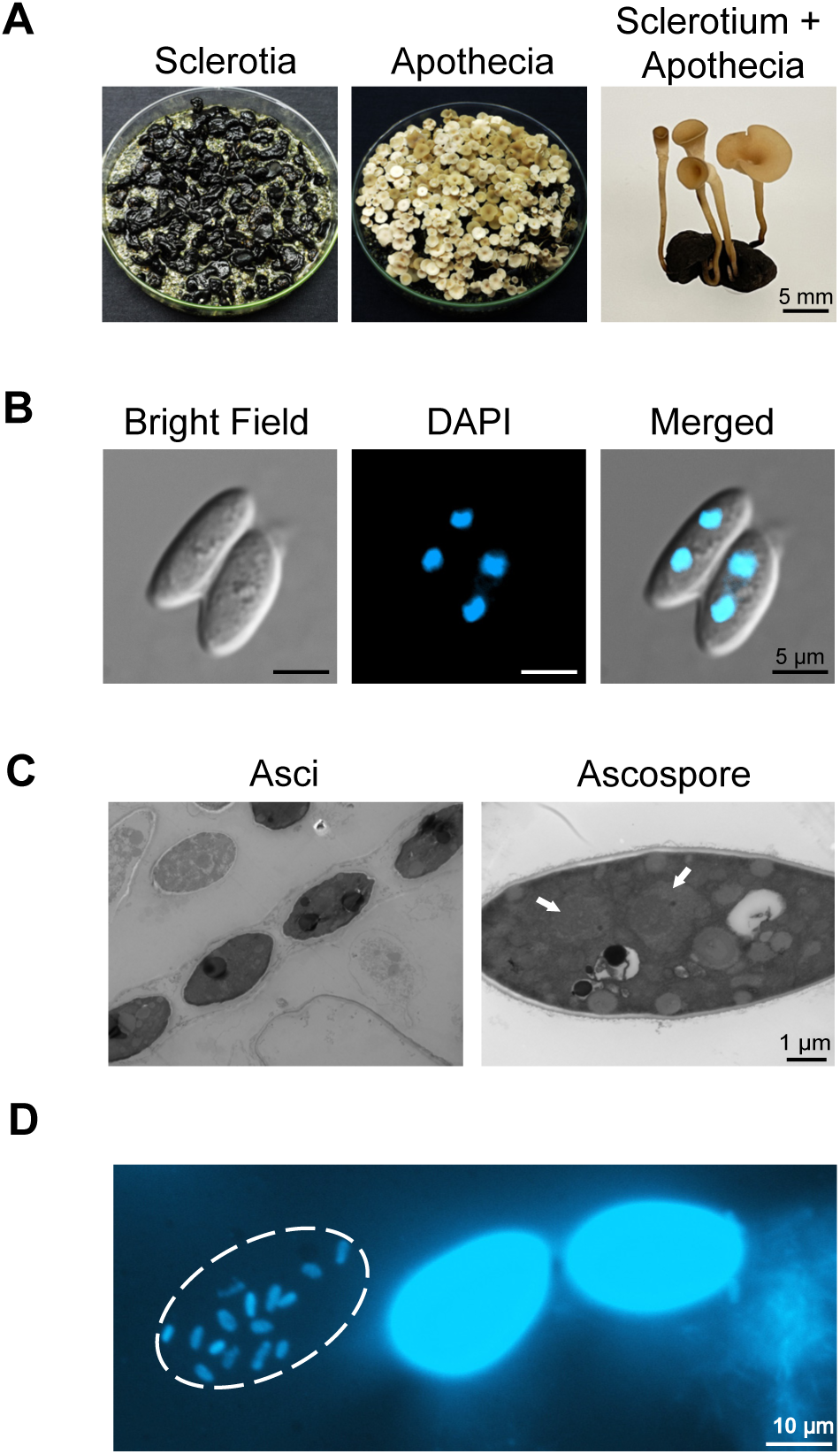
*S. sclerotiorum* ascospore contains two nuclei, each with mostly 8 chromosomes. A, The formation of apothecia from sclerotia. Scale bar: 5 mm. B, DAPI staining of ascospores. Blue signals represent the nuclei. Scale bar: 5 μm. C, TEM image of *S. sclerotiorum* ascospore. White arrows indicate the nucleolus inside the nuclei. Scale bar: 1 μm. D, DAPI staining of chromosomes released from broken ascospore. White dashed oval outlines the broken ascospore. The nearby unbroken ascospores are shown as controls. Scale bar: 10 μm.

## Molecular evidence of the presence of only one full chromosome complement inside each ascospore

Our whole genome sequencing of mutant mycelia originated from single ascospores alluded to the haploid nature of the ascospore (Supplementary Fig. S1). To confirm that each ascospore indeed contains only one set of haploid chromosomes instead of two, we first carried out fluorescence in situ hybridization (FISH) analysis using an 18S-rDNA probe from chromosome 7 (Supplementary Fig. S2). As shown in Fig. 2A and Fig. 2C, FISH hybridization signals were only observed in one nucleus per ascospore. In no case did we observed 18S-rDNA signals in both nuclei from a single ascospore, further supporting our hypothesis that the chromosome sets inside the two ascospore nuclei are different. To rule out the possibility that the probe enters only one nucleus, a control probe which can hybridize to telomeres of almost all the chromosomes (Supplementary Fig. S2) were tested. As shown in Fig. 2B-C, telomere probe signals can be seen mostly in both nuclei, supporting that the fluorescent probes can enter both nuclei freely. Taken together, these data suggest that the sets of chromosomes of the two nuclei in *S. sclerotiorum* ascospore are distinct and raising the possibility that each nucleus carries a fixed set of 8 chromosomes.

**Figure 2.**
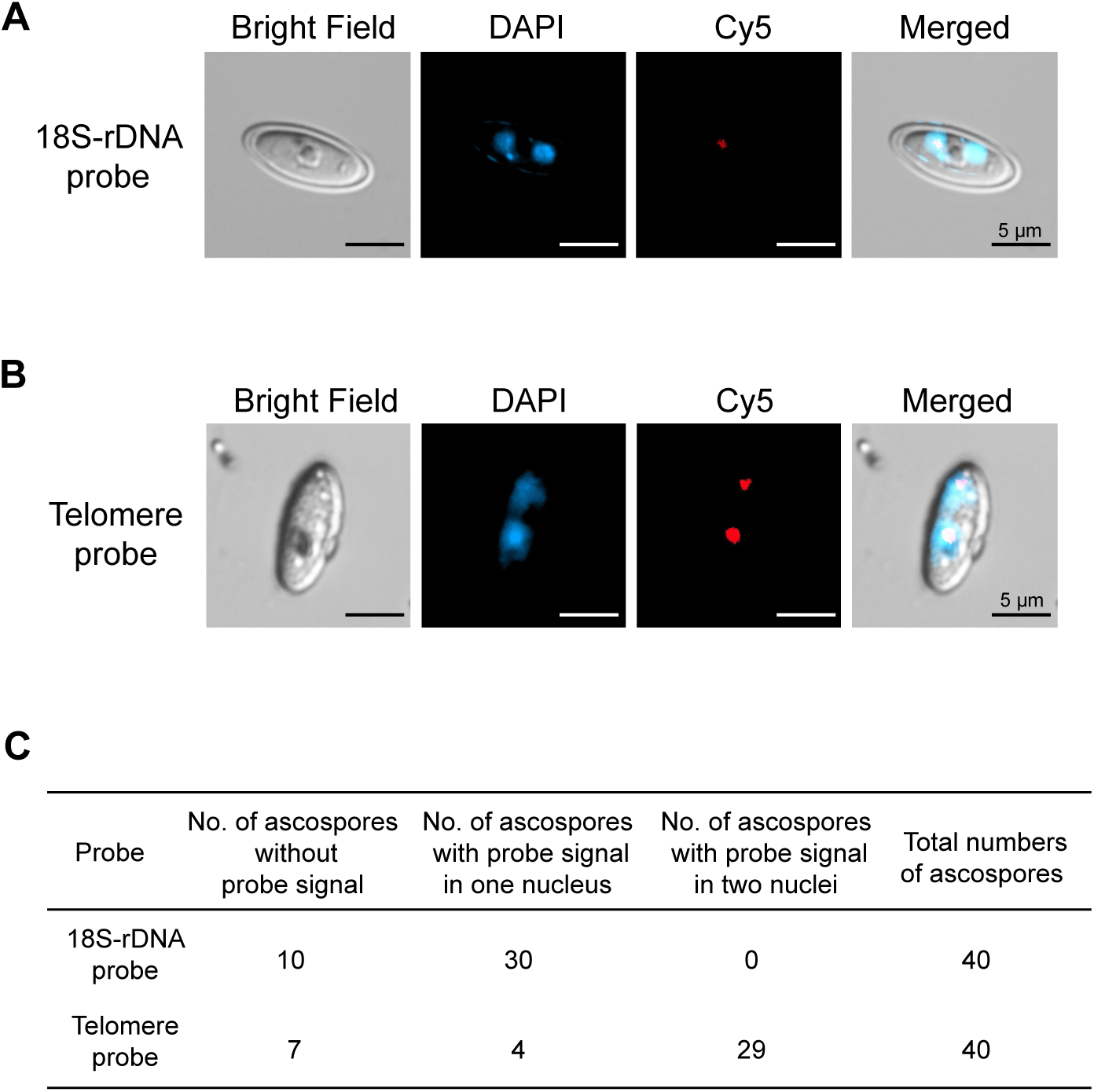
FISH analysis of ascospores of *S. sclerotiorum*. A-B, FISH signals of 18S-rDNA (A) and telomere probes (B) in asco-spores. Blue and red signals represent nuclei and probes, respec-tively. Scale bar: 5 μm. C, The numbers of ascospores show 18S-rDNA or telomere probes hybridization in one nucleus, two nuclei or no probe hybridization.

## Molecular evidence of random assortment of the 16 chromosomes into two nuclei in *S. sclerotiorum*

Our FISH data suggest it is possible that each nucleus carries fixed sets of 8 chromosomes. Although unlikely, we could not exclude that each nucleus contains a random subset of 8 chromosomes that together form the full chromosome complement. To measure the composition of chromosomes residing in each nucleus we attempted to isolate single nuclei from ascospores. However, our attempts failed due to the fragility of *S. sclerotiorum* ascospore nuclei which ruptured during nuclei extraction. We next used young mycelia for the nuclei extraction since they are the haploid descendants of an ascospore. After sucrose gradient centrifugation and fractionation (Supplementary Fig. S3A-B), the purified nuclei were singly sorted into 96-well plates using a Beckman Coulter Moflo Astrios cell sorter. After whole genome DNA amplification, DNA from each nucleus was subjected to PCR with primers specific to each chromosome. Interestingly, for most single nuclear genomes, only 8 chromosomes were present (Supplementary Fig. S4, Fig. 3A). Such result is consistent with our cell biology data, confirming that each nucleus in ascospores and young mycelium cells indeed contains about 8 chromosomes. Surprisingly, the constitution of these 8 chromosomes varied among nuclei (Supplementary Fig. S4, Fig. 3A), indicating a likely random assortment of the 16 chromosomes into different nuclei in *S. sclerotiorum* (see model in Figure 3B).

**Figure 3.**
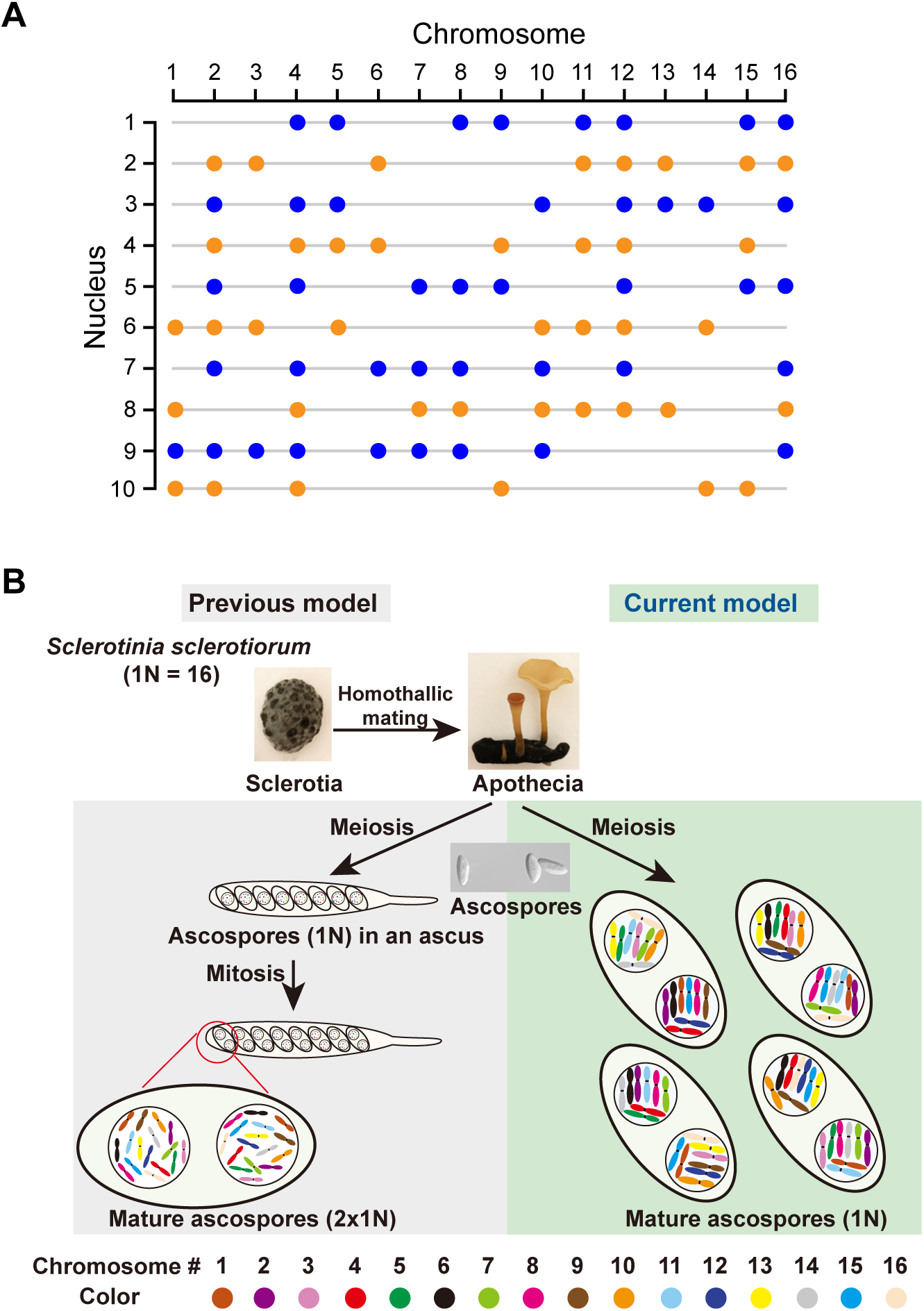
*S. sclerotiorum* sorts its full haploid chromosomes randomly into two nuclei. A, Summary table for the PCR results with chromosome-specific primers. DNA from singly sorted nucleus was subjected to whole genome amplification, and the amplified genomic DNA was used as template for PCR with chromosome-specific primers. Blue and orange dots indicate the presence of corresponding chromosomes in each nucleus. B, Models comparing the status of chromosomes in *S. sclerotiorum* nuclei. In the old model, the two nuclei in each ascospore contain the same complement of 1N chromosomes (1N=16). In the new model, the 16 chromosomes are split randomly into two nuclei.

## Random assortment of the haploid 18 chromosomes into multiple nuclei in *B. cinerea*

After observing the random assortment of 16 chromosomes of *S. sclerotiorum* into two nuclei, we asked whether such phenomenon is present in other fungi. In *B. cinerea*, each haploid conidium is also multinucleated, with predominantly 4-5 nuclei each (Shirane, Masuko and Hayashi, 1988). Its haploid genome has 18 chromosomes (1N=18) based on *de novo* genome sequencing and assembly. Using the same single nuclei purification approach (Supplementary Fig. S5A), we sorted single purified nuclei from conidia of *B. cinerea* into 96-well plates (Supplementary Fig. S5B) and analyzed for the presence of individual chromosomes by PCR. As shown in Supplementary Fig. S6 and Fig. 4A, each nucleus harbored between 3 to 7 chromosomes in a random fashion. Thus, like *S. sclerotiorum*, *B. cinerea* carries the full chromosome complement in each conidium, with the chromosome content randomly distributed across nuclei within the conidium (see model in Figure 4B). Each conidium is truly haploid. This might explain why it is much easier to obtain immediately pure knockout or transgenic colonies using *B. cinerea* conidia than *S. sclerotiorum* mycelial cells (Reis, Pfiff and Hahn, 2005).

**Figure 4.**
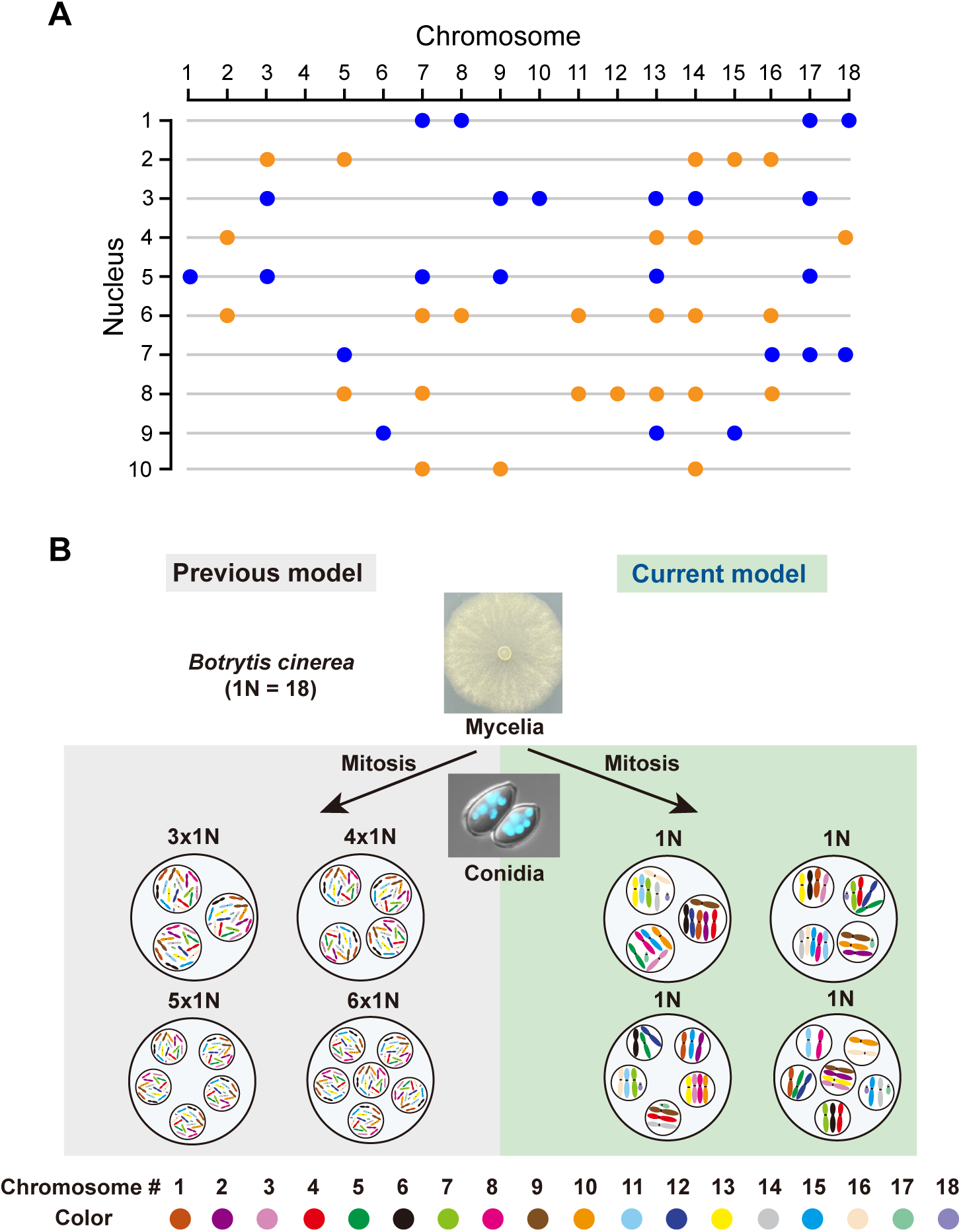
*B. cinerea* sorts its haploid chromosomes randomly into different nuclei. A, Summary table for the PCR results with chromosome-specific primers. DNA from singly sorted nucleus was subjected to whole genome amplification, and the amplified genomic DNA was used as template for PCR with chromosome-specific primers. Blue and orange dots indicate the presence of corresponding chromosomes in each nucleus. B, Models comparing the status of chromosomes in *B. cinerea* nuclei. In the old model, the nuclei in each conidia spore contain the same complement of 1N chromosomes (1N=18). In the new model, the 18 chromosomes are split randomly into different nuclei.

## Discussion

From our genetic analysis of *S. sclerotiorum* UV mutants, FISH and single nuclei sorting and PCR analysis we conclude that in both *S. sclerotiorum* and *B. cinerea* haploid chromosomes are randomly partitioned into separate nuclei. During mitosis and meiosis, how the chromosomes are segregated randomly to different nuclei while maintaining their genetic integrity is unclear. On surface, such mechanism seems chaotic. Close examination of these processes using cell biology tools will be needed to address these questions.

Why *S. sclerotiorum* and *B. cinerea* have such remarkable pattern of chromosome sorting is also unclear. One straight forward benefit may be to enable further separation of transcriptional processes of each nucleus. However, the seemingly random pattern of the assortment is puzzling. It is also unclear how wide spread such phenomenon is in fungi or even other eukaryotes. More chromosomal analyses in different non-model fungal species are needed. Future in-depth cellular and molecular genetic analysis of the genes involved in this process will reveal the intricate mechanistic details of such chromosome sorting and regulation. Understanding these mechanisms may lead to dramatic ramifications in synthetic biology and genetic engineering.

Our study has the potential to fundamentally change our view of how multinucleated cells manage their chromosomal content. Mycelial cells of many fungi could conceivably harbor hundreds of nuclei with different chromosomes, which may enable the lengthy cells to respond quickly at local foci against environmental cues. Our discovery will also change the paradigm of forward and reverse genetic analyses of fungi with multinucleate spores. For example, forward genetics is rarely attempted in species like *S. sclerotiorum* and *B. cinerea* due to the multinucleate nature of their spores. The discovery that each conidium contains only one complement of haploid chromosomes indicates that forward genetic screens can also be carried out in *B. cinerea* through random mutagenesis of conidia. Knockout experiments can also be simplified by using cell types with just one full complement of chromosomes, such as ascospores or conidia.

## Methods

### Fungal growth, apothecia induction and the collection of ascospores

*S. sclerotiorum* strain 1980 was kindly shared by Dr. Jeffrey Rollins from University of Florida. Sclerotia were collected and used to generate apothecia as described with slight modifications (Athukorala *et al*., 2010). Briefly, fresh mycelia grown on Potato Dextrose Agar (PDA) (25 g PDA powder and 15 g Agar dissolved in 1 L distilled water, autoclave at 121 for 20 min) were inoculated on carrot medium (fresh carrot was cut into 3-5 mm thick slices and placed into a sterilized glass container, autoclave at 121 for 20 min) and grown for 3-4 weeks to produce large sclerotia. The resulting sclerotia were gently brushed with test tube brush to remove the surface mycelia and carrot residues. Sclerotia were then dried at room temperature for 2 weeks. The dried sclerotia were surface sterilized with 33% (v/v) bleach solution (Clorox Disinfecting Bleach) for 5-10 min and washed with sterile water twice. The clean sclerotia were placed on autoclaved sand in glass petri dishes (14 cm diameter). The petri dishes were incubated at 4 for 2-4 weeks before being transferred to a 16 -plant growth incubator under 16 h light/8 h darkness cycles. The formation of apothecia is visible in 3-5 weeks.

Ascospores were liberated by rigorously vortex of the fully open apothecia in a 2 ml Eppendorf tube with distilled water. The resulting liquid was filtered with Corning Cell Strainer (pore size 40 μm) and Whatman filter papers (90 mm diameter) to remove mycelia, tissue chunks and other contaminants. The filtrate was centrifuged at 12000 rpm for 2 min and the supernatant was discarded. The remaining pellet constitutes pure ascospores, which is confirmed with microscopy and DAPI staining.

### Observation of nuclei and chromosomes

Nuclei observation: Ascospores were treated with ethanol-acetic acid solution (ethanol:acetic acid = 3:1, v/v) for 30 min on glass slides. The specimens were dehydrated in an ethanol series of 70%, 90% and 100% at 5 min intervals, and air dried. Then the slides were stained with premixed DAPI solution (1 μg/ml) and imaged under a ZEISS fluorescence microscope.

Chromosome observation: A cell burst method was employed to observe chromosomes as described with slight modifications (Taga and Murata, 1994). In brief, ascospores were treated with methanol-acetic acid solution (methanol:acetic acid = 1:1, v/v) for 30 min on glass slides and flame dried. The specimens were transferred to 95% ethanol and kept in 70% ethanol for 3 hr. After hydrolyzation in 1N HCl for 5 min at room temperature and for 10 min at 60°C, slides were washed with distilled water and stained with premixed DAPI solution (1 μg/ml) and imaged under a ZEISS fluorescence microscope.

For observation of nuclei under transmission electron microscope (TEM), ascospores were fixed using a 4% paraformaldehyde and 2.5% glutaraldehyde mixture in 0.1M cacodylate buffer, pH 6.9-7.4. Specimens were stained using 1% osmium tetroxide for 3 hours and then dehydrated in an ethanol series of 25%, 50%, 75%, 90% and 1x acetone at 15 min intervals. Specimens were embedded in acetone/Spurr resin mixture (acetone:Spurr resin = 3:1, 1:1, 1:3) and pure resin for 2 hour intervals and then were allowed to polymerize at 60 °C for 24 hours (Spurr, 1969). Resin samples were sectioned (60-100nm) and grids were stained with 1% uranyl acetate for 6 minutes and lead citrate for 4 minutes (Reynolds, 1963). Specimens were imaged under a FEI Tecnai™ Spirit TEM (120 kV).

### Fluorescence in situ hybridization (FISH) analysis

Probe design: The 18S-rDNA and telomere probes were designed using pre-labeled oligonucleotide probes (PLOPs) as described by Waminal (Waminal *et al*., 2018). Probes with 5’ Cy5 modification were synthesized by IDT (Integrated DNA Technologies).

FISH was performed following a described procedure (Waminal *et al*., 2018). Briefly, 32 μl of FISH hybridization master mix (50% formamide, 10% dextran sulfate, and 2× SSC) and 50 ng of probe were combined, followed by the addition of distilled water to a total volume of 40 μl. Then the glass slides with the addition of hybridization mix were sealed with a glue gun to prevent evaporation. The sealed glass slides were placed at 80 for 10 min to denature genomic DNA. Then the hybridization was performed overnight at 37. The glass slides were then washed using 2× SSC at room temperature for 5 min, 0.1× SSC at 37 for 20 min, and 2× SSC for 10 min at room temperature, and then dehydrated in an ethanol series of 70 %, 90 % and 100 % at 5 min intervals, and air dried. Lastly, the slides were stained with premixed DAPI solution (1 μg/ml) and imaged under a ZEISS fluorescence microscope.

### Fluorescence microscopy

Stained ascospores, nuclei and FISH were all imaged using a ZEISS light microscope (ZEISS AXIO Imager M2) equipped with X-cite series 120Q for fluorescence illumination. DAPI, mRF12 and differential interference contrast (DIC) channels were used to visualize nuclei, Cy5 signals, and ascospores, respectively. Images were taken under a 20× or 100× magnification with the scale bars shown in each image.

### Nuclei isolation and purification

The fungal nuclei were purified as described with some modifications (Gealt, Sheir Neiss and Morris, 1976). The *B. cinerea* WT strain B05.10 was kindly shared by Dr. Amir Sharon. It was grown on GYM (10 g Malt extract, 4 g glucose, 4 g yeast extract and 15 g agar dissolved in 1 L water, pH 5.5) plate. Cultures were incubated at 21 under regular florescent light for around 10 days with 16 h light/8 h darkness for conidia induction. To harvest the conidia, a sterile loop or spatula were used to scrape the spores off the surface of the media. The mixture was transferred to a sterile 50 ml screw cap conical tube and stored at −80 before use.

100 mg of frozen pellets of conidia or young mycelia were transferred into pre-cooled mortars and grinded into powder using liquid nitrogen, respectively. When 25 to 50 % of cells were broken, which was guided through microscopic examination, the powder was transferred into new conical tubes and resuspended with 10 ml homogenization buffer [10 mM PIPEs (Fisher Scientific), 5 mM CaCl_2_, 5 mM MgSO_4_ and 0.5 M sucrose]. All following manipulations and centrifugations were carried out at 4. The resulting homogenate was filtered through Corning Whatman filter papers (90 mm diameter) and Cell Strainer (pore size 40 μm). The filtrate was centrifuged at 6000 g for 20 min to obtain nuclei pellet. The pellet was then resuspended in 5 ml homogenization buffer and centrifuged at 100 g for 5 min to remove large mycelial pieces or other large particles passing through the previous filtrations. The supernatant was transferred into a new tube and centrifuged at 6000 g again for 20 min. The resulting pellet was resuspended in 3 ml 2.1 M sucrose buffer (10 mM PIPEs, 5 mM CaCl_2_, 5 mM MgSO_4_ and 2.1 M sucrose) and centrifuged at 23000 rpm (Beckman Ultracentrifuge, TLA-100.3 Rotor) for 5 min to remove small mycelial pieces and remaining intact cells. The resulting supernatant was then centrifuged at 55000 rpm for 1 h to obtain pure nuclei. The isolated nuclei were examined for purity under microscope and stored in Nuclei storage buffer (NSB, 20 mM Tris-HCl, pH 8.0, 75 mM NaCl, 0.5 mM EDTA, 50% (v/v) glycerol, 1 mM DTT, and 0.1 mM PMSF) at −80 (Ling and Waxman, 2013).

### Single nuclei sorting

The isolated pure nuclei were stained with 1 μg/ml DAPI (DAPI : nuclei solution = 1:10, v/v) for 10 min on ice and were then sorted into 96-well plates using a Moflo Astrios high-speed cell sorter (Beckman Coulter). Sorting was only conducted upon samples passing quality control measurements. The sorter was equipped with a standard 100 μm nozzle. The sheath pressure was kept at 20 PSI and gating was based on FSC-W (Forward Scatter Width), SSC-A (Side Scatter Area), 405-448/59-A (fluorescence emission at 448 nm with the excitation at 405 nm), which select nuclei based on their size, granularity, and fluorescence characteristics, respectively. The 96-well plates with sorted nuclei were immersed into dry ice-ethanol bath immediately for 1 min before being transferred to −80 for long-term storage.

### Whole genome amplification of single nuclei and PCR analysis

After successful single nuclei sorting into 96-well plates, each *S. sclerotiorum* or *B. cinerea* single nucleus was subjected to whole genome amplification. All following experiments were conducted in a Laminar flow hood with filter tips to prevent DNA contamination. The method was modified from a previous described protocol (Martinez-Hernandez *et al*., 2017). In brief, a 20 μl reaction mix was prepared for each nucleus, which contained 0.5 μl of phi29 DNA polymerase (Catalog # M0269L; New England Biolab), 2 μl of phi29 10 × reaction buffer (Catalog # M0269L; New England Biolab), 2 μl of 20 × exo-resistant random primers (Catalog # SO181; Thermo Fisher Scientific), 1.5 μl of 2.5 mM dNTPs (Catalog # N0447L, New England Biolab) and 14 μl autoclaved mili-Q water. The reaction mixture was incubated at 30°C for 16 h for genome amplification. After that, the samples were treated at 65°C for 10 min to deactivate the enzyme.

The amplified genome from each nucleus was subsequently used as DNA templates for PCR reactions with chromosome-specific primers. NSB after whole genome amplification was used as a negative control. The amplified genome from the singly sorted protoplast from young mycelia of *S. sclerotiorum* or *B. cinerea* conidia was used as a positive control. The primers used are listed in Supplementary Table 1 and 2. Two sets of primers were designed from two single-copy genes located distantly on each chromosome, amplifying fragments of almost identical lengths.

For each chromosome-specific PCR reaction, these primers were mixed together to amplify both genes simultaneously. 16 or 18 primer mixtures for *S. sclerotiorum* and *B. cinerea*, respectively, were prepared with each amplified single-nucleus genome as templates. The resulting PCR products were subjected to electrophoresis followed by ethidium bromide staining to examine the presence of the corresponding chromosomes.

## Acknowledgements

The authors cordially thank Dr. Jeffrey Rollins from University of Florida for *S. sclerotiorum* strain 1980, and insightful discussions on many aspects of Sclerotinia biology. Dr. Amir Sharon from Tel Aviv University is sincerely thanked for *B. cinerea* strain B05.10. Justin Wong and Andrew Johnson at the UBC Flow Cytometry (ubcFLOW) facility are sincerely thanked for their patience and assistance in our many attempts of single nuclei sorting. Dr. Junxing Lu, Dr. Weijie Huang and Yihan Gong are thanked for providing apothecia for some of the experiments. This study was financially supported by grants to XL and YZ from the Canadian Natural Sciences and Engineering Research Council (NSERC) Discovery program, NSERC-CREATE-PRoTECT, Canada Research Chair (CRC), and the Canadian Foundation for Innovation (CFI) funds. Scholarships to YX and LT were from the Chinese Scholarship Council, and scholarship to JT was from the University of British Columbia Four-year fellowship program.

## Author Contributions

XL and YZ conceived the idea. XL, YZ and NO’N designed the experiments. YX, LT, JT and JL carried out the experiments. All authors analyzed data. XL and YZ supervised the work. YX, LT and XL wrote the draft manuscript, and all authors contributed to the revisions.

**Correspondence and requests for materials** should be addressed to XL.

---

**Supplementary Figure S1.**
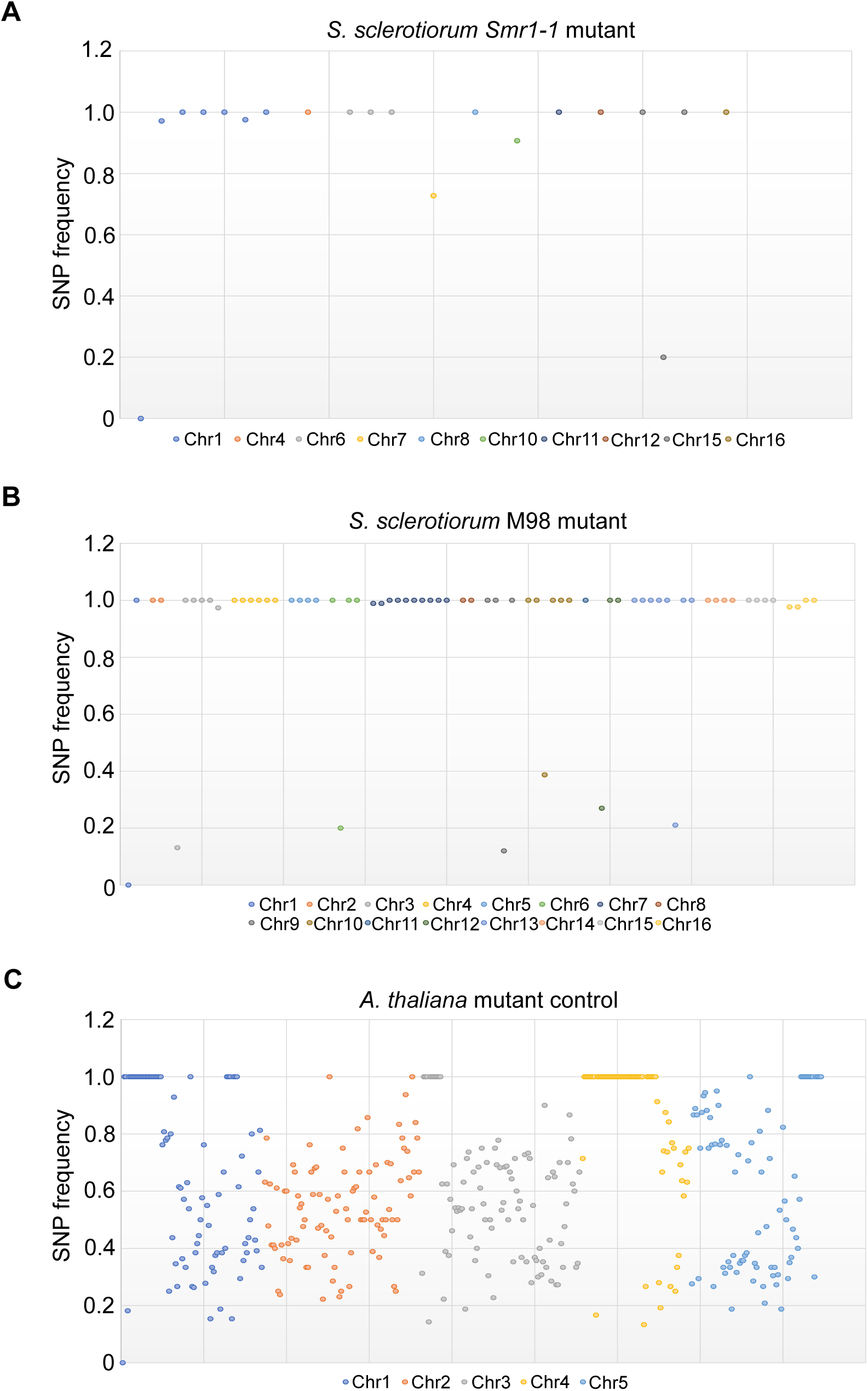
The SNP/Indel frequencies in mutants that underwent next-generation sequencing from *S. sclerotiorum* and *Arabidopsis thaliana*. A-C, Dot plot of frequencies of single nucleotide polymorphism (SNP) and inser-tion deletion (Indel) mutations on specific chromosomes originated from *Sssmr1-1* (A) and M98 (B) UV mutants in *S. sclerotiorum* (Xu *et al*., 2022), and 116-6 (C) in *A. thaliana* (SNP data from an unpublished EMS mutant, serving as a diploid mutant control). SNPs located within repetitive gene clusters and SNPs shared among multiple independent mutants are filtered out as they are likely sequencing errors.

**Supplementary Figure S2.**
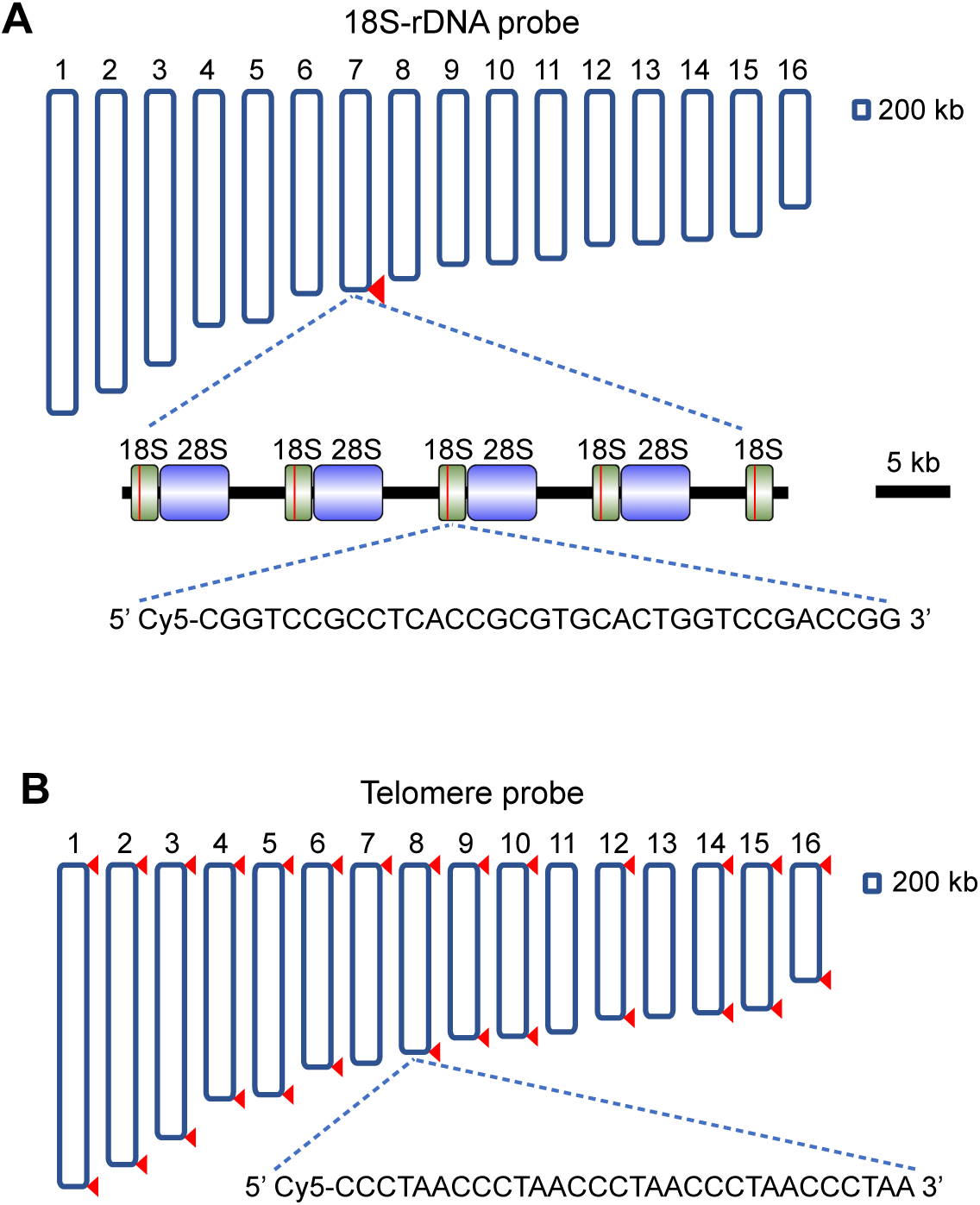
Description of the 18S rDNA and telomere probes. A and B, the diagrams of 18S rDNA and telomere probes. The positions where probes can hybridize to chromosomes of *S. sclerotiorum* are indicated by red triangles. The five 18S-rDNA probe regions are indicated by red lines. The sequences of probes are shown at the bottom of each panel. The scale bars showing the lengths of chromosomes and rDNA tandem repeats are labelled on the right.

**Supplementary Figure S3.**
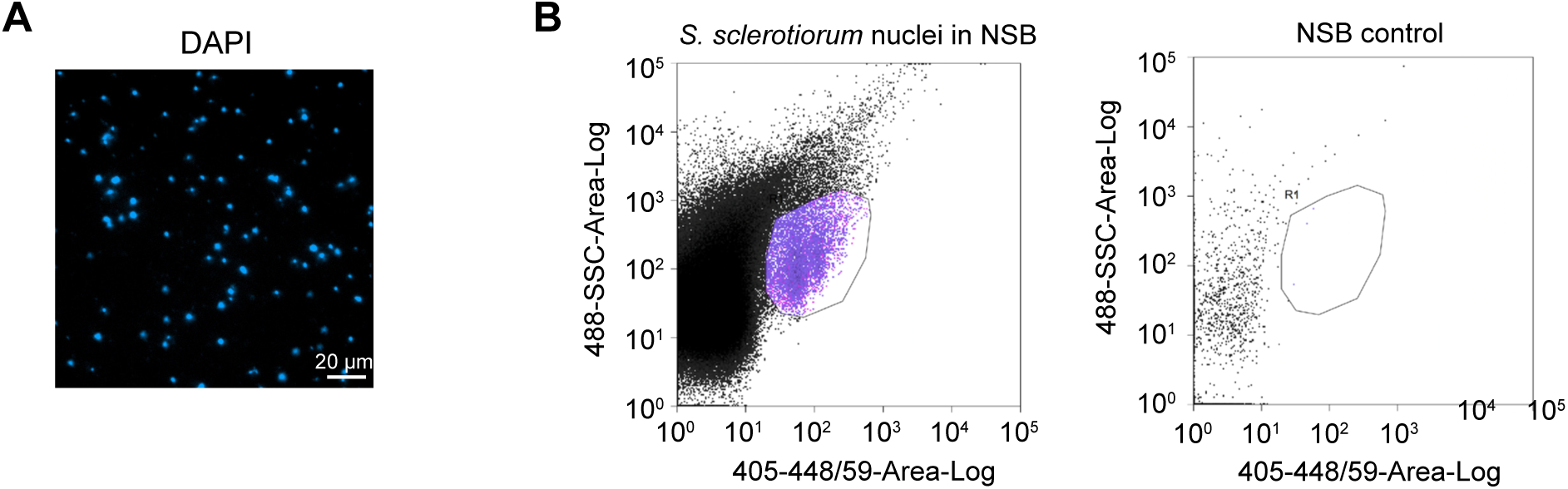
The isolation and sorting of nuclei of S. sclerotiorum. A, Purified nuclei of *S. sclerotiorum* after sucrose gradient ultracen-trifugation. Scale bar: 20 μm. B, Quality check from the Beckman cell sorter. The circled region with DAPI signals was selected for sorting. NSB (nuclei storage buffer) was used as negative control.

**Supplementary Figure S4.**
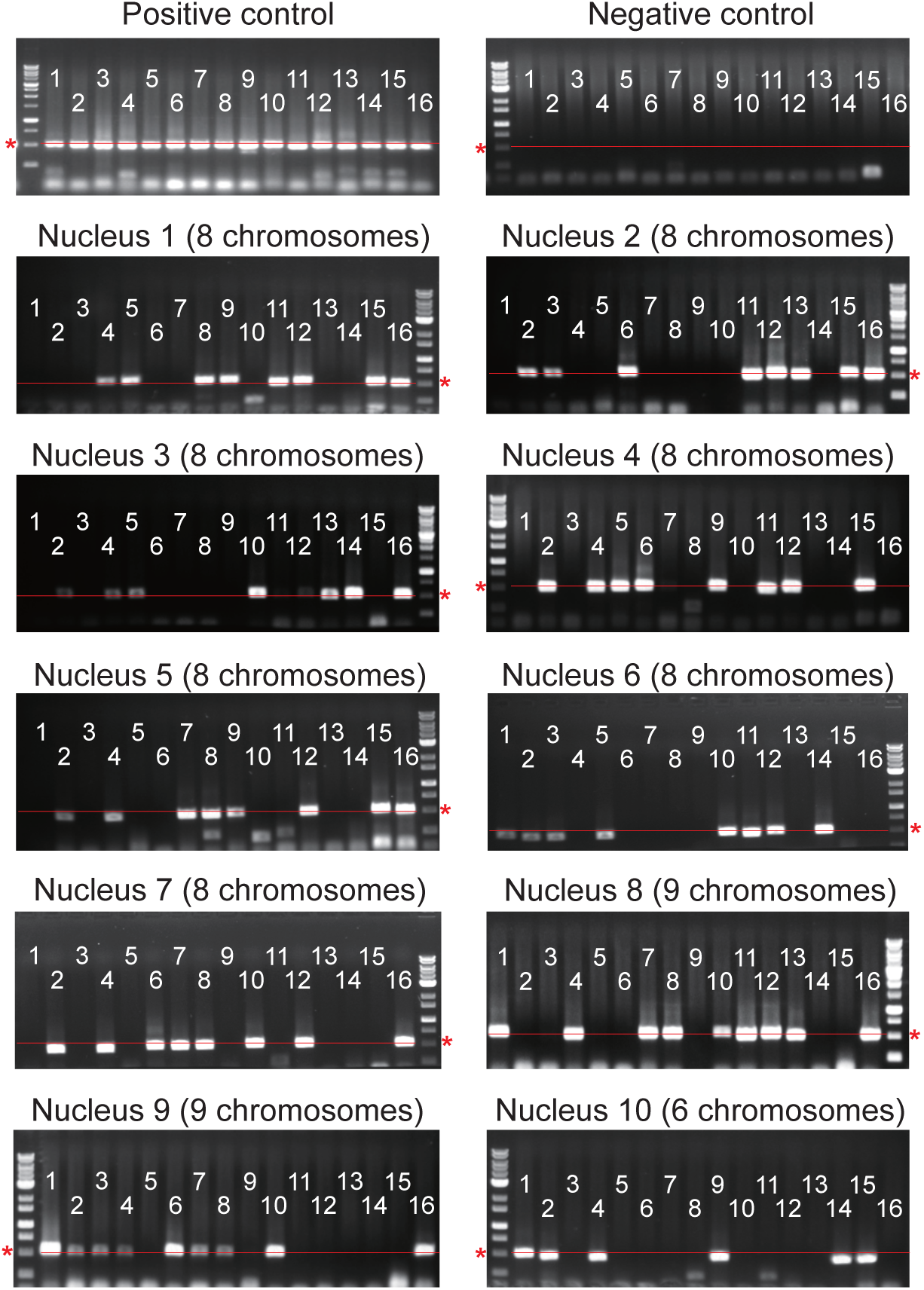
Representative gel images of PCR results after gel electrophoresis for *S. sclerotiorum* (summarized in. **Fig. 3A). Asterisks and red lines indicate the position of 500 bp.**

**Supplementary Figure S5.**
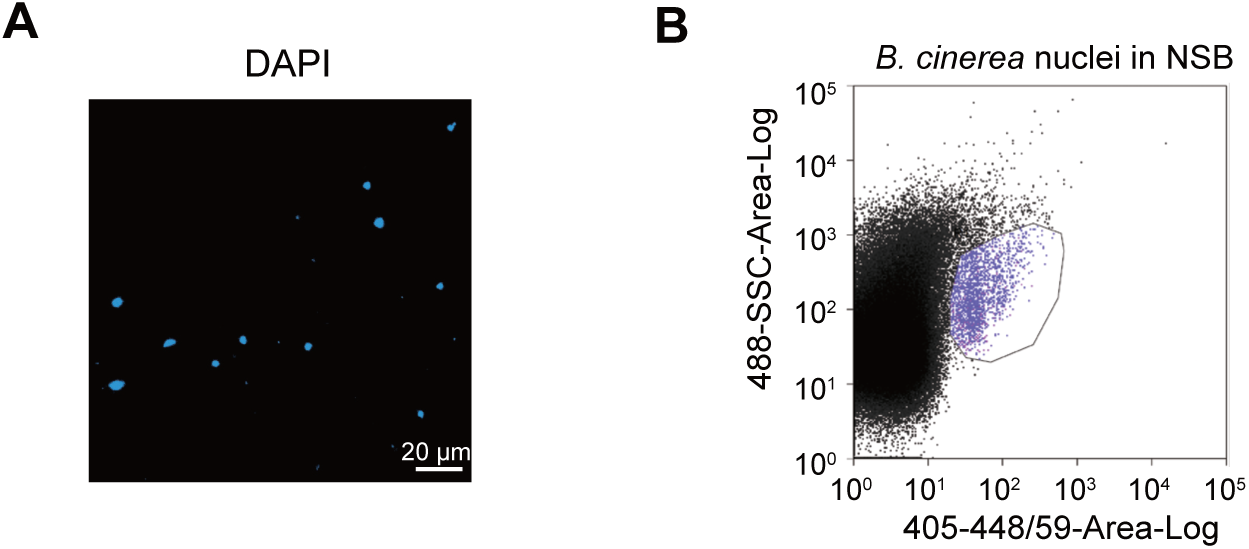
The isolation and sorting of nuclei of B. cinerea. A, Purified nuclei of *B. cinerea* conidia after sucrose gradient ultracentrifugation. B, Quality check from the Beckman cell sorter. The circled region with DAPI signals was selected for sorting. The negative control was the same as Fig. S3B.

**Supplementary Figure S6.**
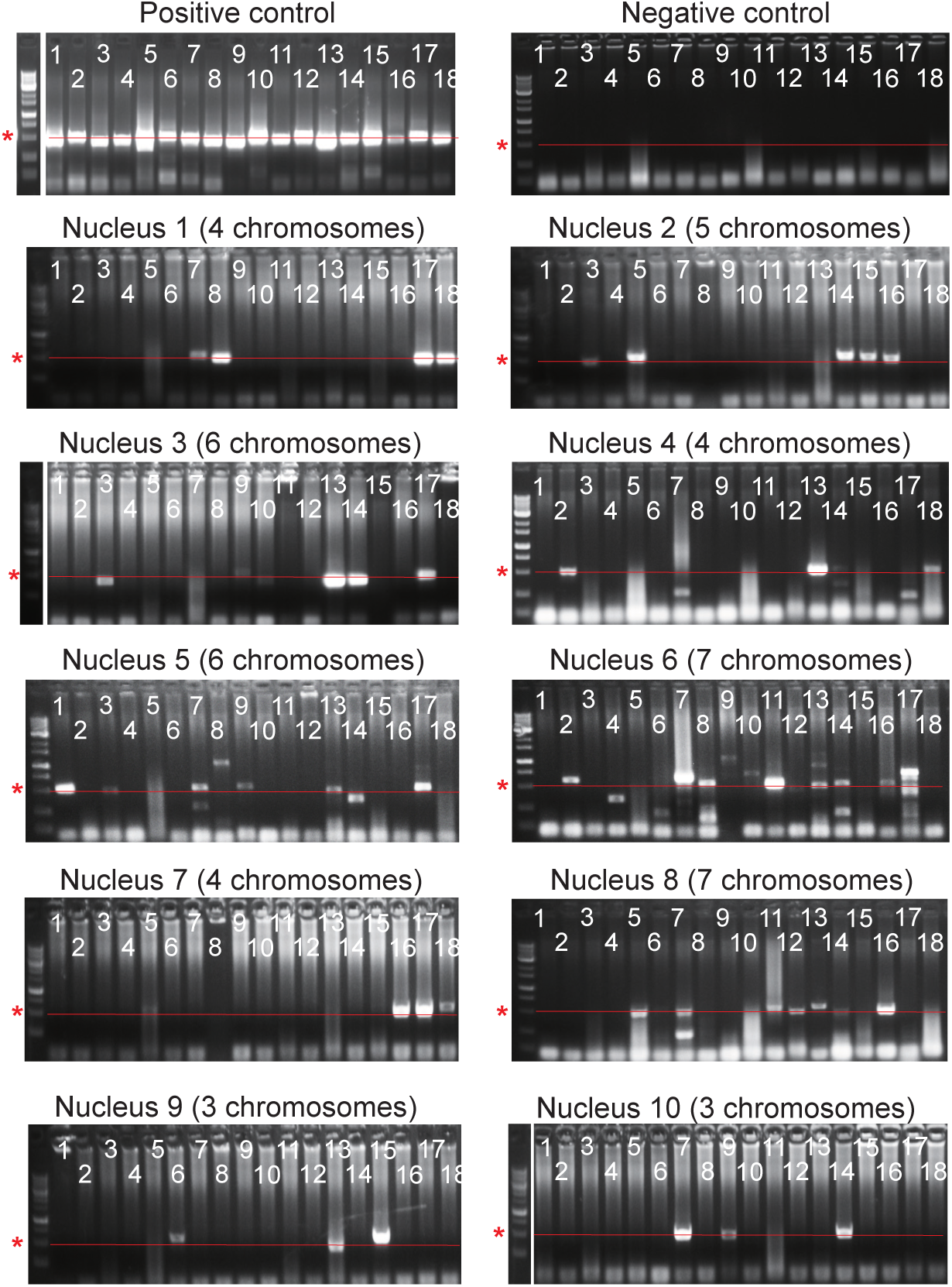
Representative gel images of PCR results after gel electrophoresis for *B. cinerea* (summarized in Fig. 4A). Asterisks and red lines indicate the position of 500 bp.

**Supplementary Table S1:**
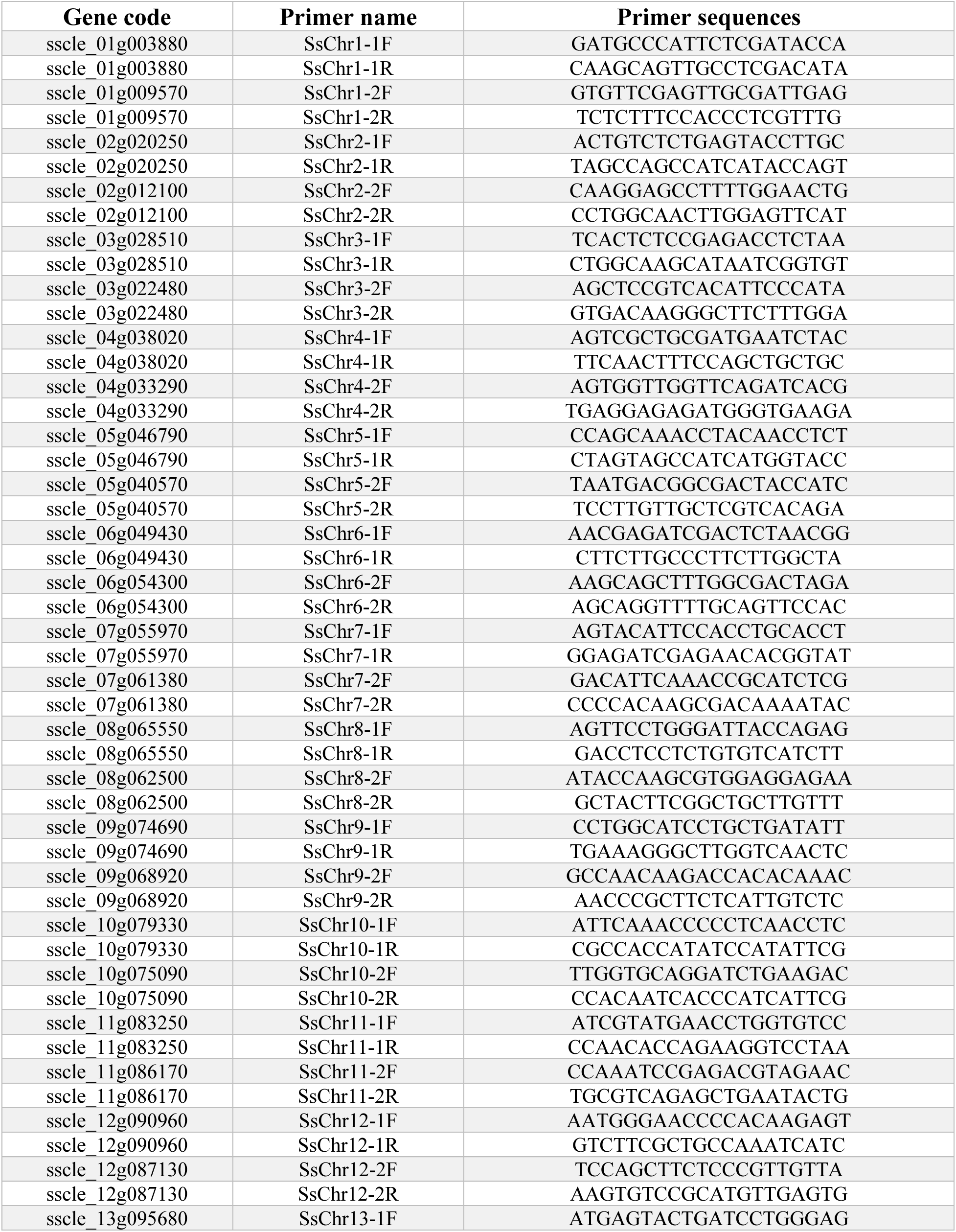
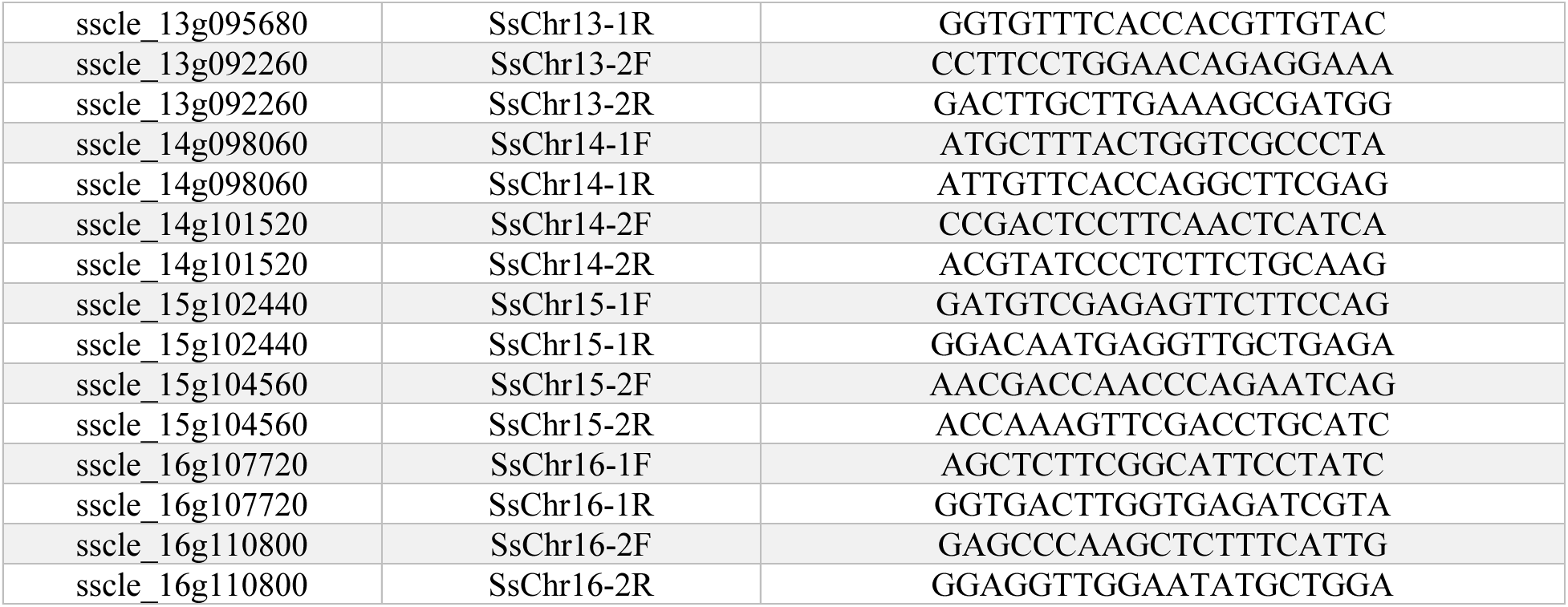
The summary of primers specific to individual chromosomes of *S. sclerotiorum*.

**Supplementary Table S2:**
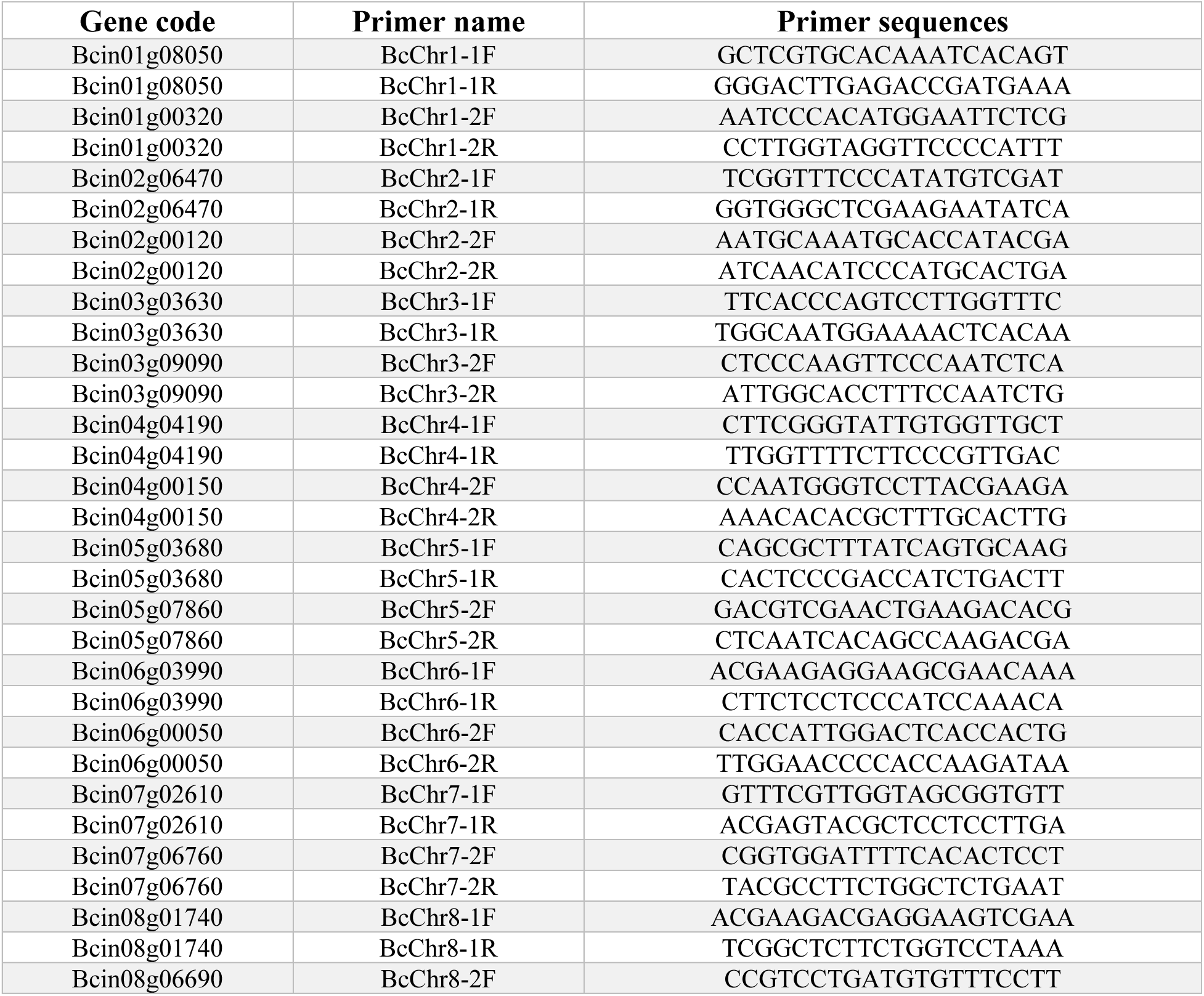
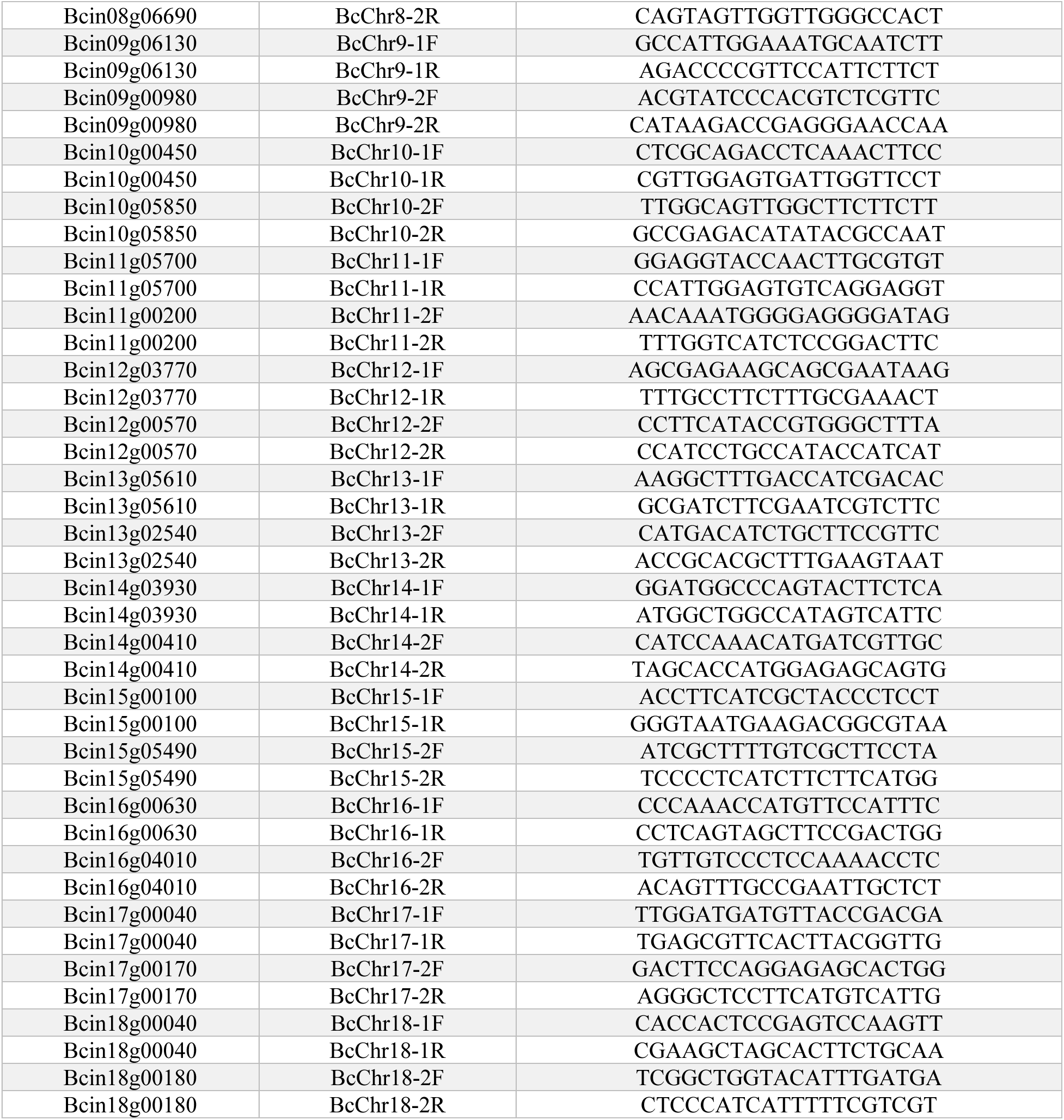
The summary of primers specific to individual chromosomes of *B. cinerea*.

## References

Adams, P. B. and Ayers, W. A. (1979) ‘Ecology of Sclerotinia species’, Phytopathology, 69(8), pp. 896–898.

Amselem, J. et al. (2011) ‘Genomic analysis of the necrotrophic fungal pathogens Sclerotinia sclerotiorum and Botrytis cinerea’, PLoS genetics, 7(8), p. e1002230.

Athukorala, S. N. P. et al. (2010) ‘The role of volatile and non-volatile antibiotics produced by Pseudomonas chlororaphis strain PA23 in its root colonization and control of Sclerotinia sclerotiorum’, Biocontrol Science and Technology, 20(8), pp. 875–890. doi: 10.1080/09583157.2010.484484.

Bolton, M. D., Thomma, B. P. and Nelson, B. D. (2006) ‘Sclerotinia sclerotiorum (Lib.) de Bary: biology and molecular traits of a cosmopolitan pathogen’, Molecular plant pathology, 7(1), pp. 1–16. doi: 10.1111/j.1364-3703.2005.00316.x.

Ekins, M. (1999) Genetic Diversity in Sclerotinia species.

Ford, E. J. et al. (1995) ‘Heterokaryon formation and vegetative compatibility in Sclerotinia sclerotiorum’, Mycological Research, 99(2), pp. 241–247. doi: 10.1016/S0953-7562(09)80893-9.

Fraissinet-Tachet, L., Reymond-Cotton, P. and Fèvre, M. (1996) ‘Molecular karyotype of the phytopathogenic fungus Sclerotinia sclerotiorum’, Current Genetics, 29(5), pp. 496–501. doi: 10.1007/BF02221520.

Gealt, M. A., Sheir Neiss, G. and Morris, N. R. (1976) ‘The isolation of nuclei from the filamentous fungus Aspergillus nidulans’, Journal of General Microbiology, 94(1), pp. 204–210. doi: 10.1099/00221287-94-1-204.

Hegedus, D. D. and Rimmer, S. R. (2005) ‘Sclerotinia sclerotiorum: when “to be or not to be” a pathogen?’, FEMS Microbiology Letters, 251(2), pp. 177–184. doi: 10.1016/j.femsle.2005.07.040.

Liang, X. and Rollins, J. A. (2018) ‘Mechanisms of broad host range necrotrophic pathogenesis in Sclerotinia sclerotiorum’, Phytopathology, 108(10), pp. 1128–1140. doi: 10.1094/PHYTO-06-18-0197-RVW.

Ling, G. and Waxman, D. J. (2013) ‘Isolation of nuclei for use in genome-wide DNase hypersensitivity assays to probe chromatin structure’, Methods in Molecular Biology, 977(7), pp. 13–19. doi: 10.1007/978-1-62703-284-1_2.

Martinez-Hernandez, F. et al. (2017) ‘Single-virus genomics reveals hidden cosmopolitan and abundant viruses’, Nature Communications, 8(May). doi: 10.1038/ncomms15892.

Reis, H., Pfiff, S. and Hahn, M. (2005) ‘Molecular and functional characterization of a secreted lipase from Botrytis cinerea’, Molecular Plant Pathology, 6(3), pp. 257–267. doi: 10.1111/j.1364-3703.2005.00280.x.

Reynolds, E. S. (1963) ‘The use of lead citrate at high pH as an electron-opaque stain in electron microscopy’, J Cell Biol. 17(1), pp. 208–212. doi: 10.1083/jcb.17.1.208.

Shirane, N., Masuko, M. and Hayashi, Y. (1988) ‘Nuclear behavior and division in germinating conidia of Botrytis cinerea’, Phytopathology, 78, pp. 1627–1630.

Spurr, A. R. (1969) ‘A low-viscosity epoxy resin embedding medium for electron microscopy’, Journal of Ultrastructure Research, 26, pp. 31–43. doi.org/10.1016/S0022-5320(69)90033-1

Taga, M. and Murata, M. (1994) ‘Visualization of mitotic chromosomes in filamentous fungi by fluorescence staining and fluorescence in situ hybridization’, Chromosoma, 103(6), pp. 408–413. doi: 10.1007/BF00362285.

Waminal, N. E. et al. (2018) ‘Rapid and efficient fish using pre-labeled oligomer probes’, Scientific Reports, 8(1), pp. 1–10. doi: 10.1038/s41598-018-26667-z.

Willetts, H. J. and Wong, J. A. (1980) ‘The biology of Sclerotinia sclerotiorum, S. trifoliorum, and S. minor with emphasis on specific nomenclature’, The Botanical Review, 46, pp. 101–165. doi: 10.1007/BF02860868

Xu, Y. et al. (2022) ‘A forward genetic screen in Sclerotinia sclerotiorum revealed the transcriptional regulation of its sclerotial melanization pathway’, Molecular Plant-Microbe Interactions, 35(3), pp. 244–256. doi: 10.1094/MPMI-10-21-0254-R.

